# Reproducibility Of Parameter Learning With Missing Observations in Naive Wnt Bayesian Network Trained on Normal/Adenomas Samples and Doxycycline Treated LS174T Cell Lines

**DOI:** 10.1101/014076

**Authors:** Shriprakash Sinha

## Abstract

**Insight, Innovation and Integration:** Doxycycline, a derivative of tetracycline, induces gene expression via reversible transcriptional activation. Levels of /3-catenin and other intra/extracellular genetic factors have been influenced in colorectal cancer cell lines, which make doxycycline a potential candidate for cancer chemotherapy. With the aim to build better computational models that show good prediction on test datasets, doxycycline treated cell lines might provide best training samples. This work tests the reproducibility of parameter learning and predictions based on the estimated parameters, using the Naive Bayesian Networks for Wnt pathway in case of missing observations for different nodes. The in silico experiments show the efficacy of causal models as one of the emerging diagnostic tools in development of targeted cancer therapy.

Recent efforts in predicting Wnt signaling activation via inference methods have helped in developing diagnostic models for therapeutic drug targeting. In this manuscript the reproducibility of parameter learning with missing observations in a Bayesian Network and its effect on prediction results for Wnt signaling activation is tested, while training the networks on doxycycline treated LS174T cell lines as well as normal and adenomas samples. This is done in order to check the effectiveness of using Bayesian Network as a tool for modeling Wnt pathway when certain observations are missing. Experimental analysis suggest that prediction results are reproducible with negligible deviations. Anomalies in estimated parameters are accounted for due to the Bayesian Network model. Also, an interesting case regarding usage of hypothesis testing came up while proving the statistical significance of different design setups of the BN model which was trained on the same data. It was found that hypothesis testing may not be the correct way to check the significance between design setups for the aforementioned case, especially when the structure of the model is same. Finally, in comparison to the biologically inspired models, the naive bayesian model may give accurate results but this accuracy comes at the cost of loss of crucial biological knowledge which might help reveal hidden relations among intra/extracellular factors affecting the Wnt pathway.

## 1 Introduction

Bayesian network (BN) being a collection of probabilistic classifiers or regressors constrained by conditional relationships Heckerman *et al*.^1^, serve as useful models for inference when data is missing or certain prior causal relations need to be incorporated. For simple models which are acyclic in nature, many inference algorithms exist that learn the parameters (here the conditional probability tables) and given an evidence, help in estimation of the uncertainty or probability regarding the feasibility of an event. Parameter learning is an important aspect when certain observations go missing and it becomes necessary to estimate the parameters in order to infer aspects about certain event in the network. In context of signaling pathways, the most recent work by Yoruk *et al*.^2^ develops statitical models for inference and establishes reporducibility via repeated sub-sampaling validation. Reproducibility of parameters via simulations help ascertain effectiveness of the models used under varying conditions were observations might be missing. This work focuses on developement of a framework to test the different design setups has been presented to investigate the reproducibility of parameters for a BN model.

Verhaegh *et al*.^3^ present an extensive study of knowledge based computational models to identify tumor driving signaling pathways using primitive designs of naive BNs that incorporate minimal biological knowledge in various cancer types. A similar but more focused work by Sinha^4^ goes further to compare biologically inspired models with a modification of the naive Bayesian Network model for Wnt pathway in human colorectal cancer case. It has been shown that the biologically inspired models reveal hidden biological relations among the intra/extracellular factors affecting the pathways at a lower level of accuracy compared the naively designed models with minimal biological knowledge that show high prediction accuracies. This drop in accuracy happens due to the complexity of the knowledge incorporated in the networks. It should be noted that lower does not necessarily mean bad results but might often present a more knowledge based belief in the interpretations of the biological results.

In this work, the issues related to use of naive BN models as knowledge based tools for identifying tumor driven signaling pathways has been explored. The manuscript tests the effectiveness of using BN as a tool for modeling Wnt pathway when certain observations are missing. Experimental analysis suggest that prediction results are reproducible with negligible deviations. Anomalies in estimated parameters are accounted for due to the BN model. Also, an interesting case regarding usage of hypothesis testing came up while proving the statistical significance of different design setups (that is scenarios of missing observations) of the BN model which was trained on the data. It was found that hypothesis testing may not be the correct way to check the significance between design setups for the aforementioned case, especially when the structure of the model is same but the nodes where the observations are missing is different.

Adopting a primitive structure of the Wnt signaling pathway from Clevers^5^, the Wnt switch being active or inactive can be encoded by investigating whether the transcription complex (*TRCMPLX*) is active or inactive. The *TRCMPLX* begins the transcription of a gene based on the activation of Wnt signaling pathway at the cell membrane. This is shown in figure 1(B). On capture of Wnt protein, the destruction complex that helps in the phosphorylation of *β*-catenin and its destruction via ubiquitination (figure 1(A)), is not formed. Due to this, *β*-catenin becomes available in an amount that is more than the required quantity. Excess *β*-catenin on reaching the nucleus helps in the formation of the *TRCMPLX*, by dislodging the Groucho (a lock that keeps the *TCF4* from helping in transcription) and combining with the TCF4. The *TRCMPLX* then begins the transcription of the Wnt target genes. These transcriptions are indicators of the cancer at various stages in different parts of the body. It is presumed Clevers^5^, that the Wnt signaling is a universal pathway that contributes in significant ways to various types of cancer, but the behaviour of the pathway itself is not fixed. Thus cancer in hair follicles or colon or skin may all have contributions from the Wnt pathway, but there will be slight/significant variations in the modus operandi of the pathway and proteins acting in it. This asks for development of diagnostic computational models which may help in predicting the state of the Wnt pathway for a specific cancer. The Wnt pathway being active or inactive can be tested by inferencing whether the transcription complex (*TRCMPLX*) is active or inactive. This is done by capturing the expression values of the Wnt target genes using a reference BN from Verhaegh *et al*. ^6^.

**Fig. 1.**
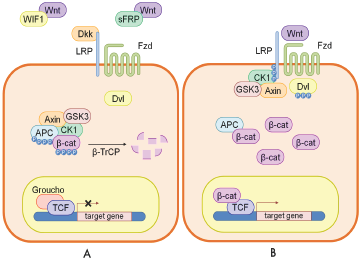
A cartoon of wnt signaling pathway adapted and redrawn from Clevers^5^ and Verhaegh *et al*.^6^. Part (A) represents the destruction of *β*-catenin leading to the inactivation of the wnt target gene. Part (B) represents activation of wnt target gene.

Tetracyclines have been extensively used in many of the cancer types and have found effective to a certain extent in chemotherapeutic treatments in colorectal cancer (Gu *et al.*^7^). Tetracycline controlled transcriptional activation is a method of inducible gene expression where transcription is reversibly turned on or off in the presence of the antibiotic tetracycline or one of its derivatives like doxycycline (Berens and Hillen^8^, Jardé *et al*.^9^ (http://en.wikipedia.org/wiki/Tetracycline-controlled_transcriptional_activation)). Current research has stressed on the use of doxycycline (with stereochemical structure in figure 2 (http://en.wikipedia.org/wiki/File:Doxycycline.svg)) either separately (Mokry *et al*.^10^, Scholer-Dahirel *et al*.^11^) or in combination with other drugs (Sagar *et al*.^12^, Lazarova *et al*.^13^) for therapeutic targeting of Wnt signaling pathway. In recent efforts towards targeting of colorectal cancer cases (Onoda *et al*.^14^, Onoda *et al*.^15^), doxycycline has been found to effect the *β*-catenin levels (Jardé *et al*.^9^) as well as some of the intra/extracellular genetic factors affecting the Wnt pathway (Scholer-Dahirel *et al*.^11^, Noubissi *et al*.^16^)

**Fig. 2.**
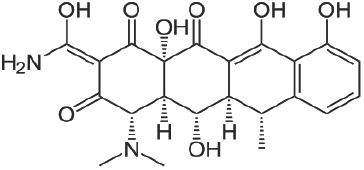
Doxycycline structure provided by Dr. Al. K. Lisch on wikipedia.

It has been found that the advantages of using doxycycline over derivatives of tetracyclines is greater and is thus a potential candidate for cancer chemotherapy (Sagar *et al*.^12^). With the aim to build better computational models that show good prediction on test datasets, doxycycline treated cell lines might provide best training samples. The network was trained on one such dataset containing measurements of expression levels for a set of Wnt target genes. Inferencing the state of the *TRCMPLX* based on the evidence provided by expression levels of the same Wnt target genes for an unlabeled sample leads to prediction of the state of Wnt pathway.

*The aim of this work is to test the reproducibility of parameter learning and predictions based on the estimated parameters, using the Wnt activation BN, in case of missing observations for different nodes*. This is done to check if the models perform well in case of missing observations, under different setups. In short, this is achieved via the following experimental setting:

- A reference BN (Ref BN) was trained on 12 colon cancer cell lines (complete real data, 6 Wnt on and 6 Wnt off)
- The parameter estimation and predictions are done using BNs from simulated data and real data (for each setup)
- Observations were sampled from:

i. colon cancer cell lines (real data) to train BN for each setup (12 observations)
ii. Ref BN to train BN for simulation for each setup. (10000 samples in each setup)
- Five setups are designed involving nodes with complete or missing observations

Significance of the results are achieved by comparing (*i*) the estimated parameters from both real observations and sampled observations for simulation, for every setup, with initially assigned parameters of the Ref BN, (*ii*) prediction results obtained using learned parameters per setup, with prediction results obtained using the assigned parameters of the Ref BN (*iii*) the estimated parameters from real observations and sampled observations for simulation, for every setup and *(iv)* prediction results obtained using learned parameters from real observations with prediction results obtained using learned parameters from sampled observations for simulation, per setup.

Section 2 gives a detailed description of the Ref BN. A set of experiments performed on the modifications of the Ref BN is explained in section 3. Next follows the results (section 4), and conclusion 5.

## 2 Method

The reference network has three main layers with nodes denoting (1) *β*-catenin, *TCF*4 and *TRCMPLX* (*β*-catenin and *TCF*4 are parents of *TRCMPLX*), (2) Wnt target genes as children of *TRCMPLX* and (3) probes as children of individual genes. The model has been adopted and redrawn in figure 3. Mathematically, the reference bnet can be defined Directed Acyclic Graph (DAG) G ≡ (V, E) defined by a set of vertices and edges. Here the nodes or vertices are V = {*β*-catenin, *TCF*4, *TRCMPLX*, *g*_1_, …, *g*_*n*_, *p*_1_, … *p*_*m*_} and the edges are E = { (*β*-catenin, *TRCMPLX*,) (*TCF*4, *TRCMPLX*),} ∪ {⋃_∀*i*_ (*TRCMPLX*, *g*_*i*_)} ∪ [⋃_∀*i*_(⋃_∀*j*_*i*__ (*g*_*i*_, *P*_*j*_*i*__)}], were *i* denotes the genes and *j*_*i*_, a corresponding set of probe(s) measuring the expression level of a particular gene *i*. The complexes, genes and the corresponding probe sets have been enlisted in table 1. Details of the network are as follows.

**Fig. 3.**
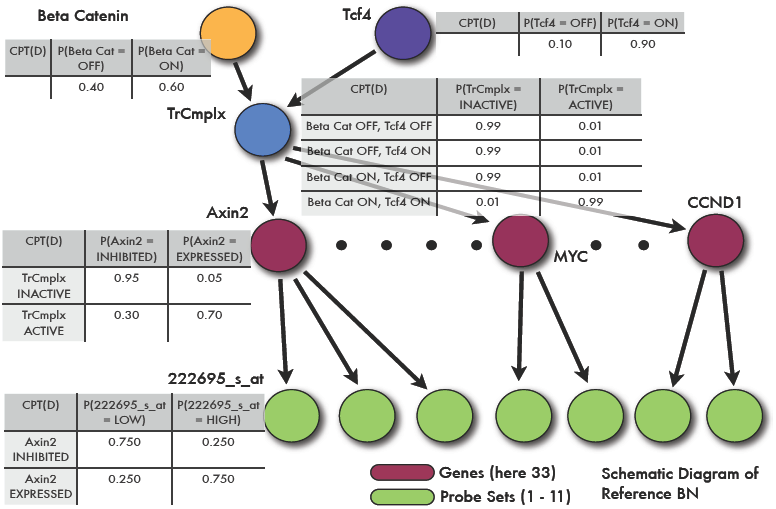
A bnet model from Verhaegh *et al*. ^6^, showing the different kinds of complexes contributing to Wnt Signaling pathway. Nodes include *β*-catenin,*TCF*4, *TRCMPLX*, the genes and their measurements in the form of probe (set) intensities.

**Table 1.** Manually fed and estimated parameter values (cpd) for nodes of a BN constructed from colon cancer cell line LS174T (Mokry *et al*. ^10^). Nodes and their parents represent the various protien complexes, genes and the corresponding probes that measure the gene expression, at different levels of the network. For colon cancer cell line data, Genes *PPARG* and *KLF*6 coloured in red indicate that they are inhibited when *TRCMPLX* is active.

| Parameter values for nodes of Bayesian net partly using Colon cancer cell lines LS174T |  |  |  |  |  |  |
| --- | --- | --- | --- | --- | --- | --- |
| No. | NODES (Parents) | Reference BNET (C)PTs | NODES (Parents) | Reference BNET (C)PTs | NODES (parents) | Reference Bnet (C)Pts |
| 1 | B-cat | [0.40 0.60] | 222695_s.at (AXIN2) | [0.75 0.25 0.25 0.75] | 221557_s.at (LEF1) | [0.625 0.375 0.375 0.675] |
| 2 | TCF4 | [0.10 0.90] | 222696.at (AXIN2) | [0.875 0.125 0.875 0.125] | 221558_s.at (LEF1) | [0.625 0.375 0.375 0.675] |
| 3 | TRCMPLX<br>(B-CAT, TCF4) | [0.99 0.99 0.99 0.01<br>0.01 0.01 0.01 0.99] | 224176_s.at (AXIN2) | [0.875 0.125 0.875 0.125] | 210515.at (HNF1A) | [0.625 0.375 0.375 0.675] |
| 4 | AXIN2 (TRCMPLX) | [0.95 0.30 0.05 0.70] | 224498_x.at (AXIN2) | [0.75 0.25 0.25 0.75] | 216930.at (HNF1A) | [0.625 0.375 0.375 0.675] |
| 5 | EPHB2 (TRCMPLX) | [0.95 0.30 0.05 0.70] | 209588.at (EPHB2) | [0.875 0.125 0.875 0.125] | 201579.at (FAT1) | [0.50 0.50 0.50 0.50] |
| 6 | EPHB3 (TRCMPLX) | [0.95 0.30 0.05 0.70] | 209589_s.at (EPHB2) | [0.875 0.125 0.875 0.125] | 226360.at (ZNF3) | [0.875 0.125 0.125 0.875] |
| 7 | MYC (TRCMPLX) | [0.95 0.30 0.05 0.70] | 210651_s.at (EPHB2) | [0.875 0.125 0.875 0.125] | 1554685_a.at (KIAA1199) | [0.875 0.125 0.125 0.875] |
| 8 | CCND1 (TRCMPLX) | [0.95 0.30 0.05 0.70] | 211165_x.at (EPHB2) | [0.875 0.125 0.875 0.125] | 212942_s.at (KIAA1199) | [0.875 0.125 0.125 0.875] |
| 9 | SP5 (TRCMPLX) | [0.95 0.30 0.05 0.70] | 1438.at (EPHB3) | [0.875 0.125 0.875 0.125] | 218704.at (RNF43) | [0.875 0.125 0.125 0.875] |
| 10 | LGR5 (TRCMPLX) | [0.95 0.30 0.05 0.70] | 204600.at (EPHB3) | [0.875 0.125 0.875 0.125] | 209081_s.at (COL18A1) | [0.50 0.50 0.50 0.50] |
| 11 | ASCL2 (TRCMPLX) | [0.95 0.30 0.05 0.70] | 202431_s.at (MYC) | [0.875 0.125 0.875 0.125] | 209082_s.at (COL18A1) | [0.50 0.50 0.50 0.50] |
| 12 | PPARG (TRCMPLX) | [0.45 0.95 0.55 0.05] | 208711_s.at (CCND1) | [0.625 0.375 0.375 0.675] | 1555832_s.at (KLF6) | [0.875 0.125 0.125 0.875] |
| 13 | CD44 (TRCMPLX) | [0.95 0.30 0.05 0.70] | 208712.at (CCND1) | [0.625 0.375 0.375 0.675] | 208960_s.at (KLF6) | [0.875 0.125 0.125 0.875] |
| 14 | SLC1A2 (TRCMPLX) | [0.95 0.30 0.05 0.70] | 235845.at (SP5) | [0.75 0.25 0.25 0.75] | 208961_s.at (KLF6) | [0.875 0.125 0.125 0.875] |
| 15 | BMP7 (TRCMPLX) | [0.95 0.30 0.05 0.70] | 210393.at (LGR5) | [0.875 0.125 0.875 0.125] | 211610.at (KLF6) | [0.875 0.125 0.125 0.875] |
| 16 | LEF1 (TRCMPLX) | [0.95 0.30 0.05 0.70] | 213880.at (LGR5) | [0.875 0.125 0.875 0.125] | 224606.at (KLF6) | [0.875 0.125 0.125 0.875] |
| 17 | HNF1A (TRCMPLX) | [0.95 0.30 0.05 0.70] | 207607.at (ASCL2) | [0.875 0.125 0.875 0.125] | 206128.at (ADRA2C) | [0.50 0.50 0.50 0.50] |
| 18 | FAT1 (TRCMPLX) | [0.95 0.30 0.05 0.70] | 229215.at (ASCL2) | [0.875 0.125 0.875 0.125] | 203705_s.at (FZD7) | [0.50 0.50 0.50 0.50] |
| 19 | ZNF3 (TRCMPLX) | [0.95 0.30 0.05 0.70] | 208510_s.at (PPARG) | [0.875 0.125 0.875 0.125] | 203706_a.at (FZD7) | [0.50 0.50 0.50 0.50] |
| 20 | KIAA1199 (TRCMPLX) | [0.95 0.30 0.05 0.70] | 1557905_s.at (CD44) | [0.75 0.25 0.25 0.75] | 202859_x.at (IL8) | [0.625 0.375 0.375 0.625] |
| 21 | RNF43 (TRCMPLX) | [0.95 0.30 0.05 0.70] | 204489_s.at (CD44) | [0.875 0.125 0.875 0.125] | 211506_s.at (IL8) | [0.625 0.375 0.375 0.625] |
| 22 | COL18A1 (TRCMPLX) | [0.95 0.30 0.05 0.70] | 204490_s.at (CD44) | [0.875 0.125 0.875 0.125] | 219682_s.at (TBX3) | [0.50 0.50 0.50 0.50] |
| 23 | KLF6 (TRCMPLX) | [0.45 0.95 0.55 0.05] | 209835_x.at (CD44) | [0.75 0.25 0.25 0.75] | 222917_s.at (TBX3) | [0.50 0.50 0.50 0.50] |
| 24 | ADRA2C (TRCMPLX) | [0.95 0.30 0.05 0.70] | 210916_s.at (CD44) | [0.75 0.25 0.25 0.75] | 225544.at (TBX3) | [0.50 0.50 0.50 0.50] |
| 25 | FZD7 (TRCMPLX) | [0.95 0.30 0.05 0.70] | 212014_x.at (CD44) | [0.75 0.25 0.25 0.75] | 229576_s.at (TBX3) | [0.50 0.50 0.50 0.50] |
| 26 | IL8 (TRCMPLX) | [0.95 0.30 0.05 0.70] | 212063.at (CD44) | [0.875 0.125 0.875 0.125] | 1553115.at (NKD1) | [0.50 0.50 0.50 0.50] |
| 27 | TBX3 (TRCMPLX) | [0.95 0.30 0.05 0.70] | 217523.at (CD44) | [0.625 0.375 0.375 0.675] | 204602.at (DKK1) | [0.50 0.50 0.50 0.50] |
| 28 | NKD1 (TRCMPLX) | [0.95 0.30 0.05 0.70] | 229221.at (CD44) | [0.50 0.50 0.50 0.50] | 207814.at (DEFA6) | [0.75 0.25 0.25 0.75] |
| 29 | DKK1 (TRCMPLX) | [0.95 0.30 0.05 0.70] | 1558009.at (SLC1A2) | [0.75 0.25 0.25 0.75] | 200648_s.at (GLUL) | [0.75 0.25 0.25 0.75] |
| 30 | DEFA6 (TRCMPLX) | [0.95 0.30 0.05 0.70] | 1558010_s.at (SLC1A2) | [0.625 0.375 0.375 0.675] | 215001_s.at (GLUL) | [0.75 0.25 0.25 0.75] |
| 31 | GLUL (TRCMPLX) | [0.95 0.30 0.05 0.70] | 208389_s.at (SLC1A2) | [0.75 0.25 0.25 0.75] | 242281.at (GLUL) | [0.625 0.375 0.375 0.625] |
| 32 | OAT (TRCMPLX) | [0.95 0.30 0.05 0.70] | 225491.at (SLC1A2) | [0.50 0.50 0.50 0.50] | 201599.at (OAT) | [0.50 0.50 0.50 0.50] |
| 33 | LECT2 (TRCMPLX) | [0.95 0.30 0.05 0.70] | 209590.at (BMP7) | [0.75 0.25 0.25 0.75] | 207409.at (LECT2) | [0.50 0.50 0.50 0.50] |
| 34 | REG1B (TRCMPLX) | [0.95 0.30 0.05 0.70] | 209591_s.at (BMP7) | [0.875 0.125 0.875 0.125] | 205886.at (REG1B) | [0.50 0.50 0.50 0.50] |
| 35 | SOX9 (TRCMPLX) | [0.95 0.30 0.05 0.70] | 211259_s.at (BMP7) | [0.75 0.25 0.25 0.75] | 202935_s.at (SOX9) | [0.75 0.25 0.25 0.75] |
| 36 | TDGF1 (TRCMPLX) | [0.95 0.30 0.05 0.70] | 211260.at (BMP7) | [0.50 0.50 0.50 0.50] | 202936_s.at (SOX9) | [0.75 0.25 0.25 0.75] |
| 37 |  |  | 210948_s.at (LEF1) | [0.625 0.375 0.375 0.675] | 206286_s.at (TDGF1) | [0.50 0.50 0.50 0.50] |

*β*-catenin and *TCF4* act as parents of *TRCMPLX*. *TRCMPLX* transcribes a set of Wnt genes that act as child nodes of the former. Parent child interaction is depicted via the directed arrows in figure 3. The measurements for each of the genes is culled via a set of probes per gene, which form the child nodes per gene. For the reference network, a parameter value per node represents the node’s (conditional) probability table. In the current scenario, the states of all the nodes are considered to be binary. This implies that *β*-catenin and *TCF*4 can be on or off. Their combination, the *TRCMPLX* can be active or inactive. The Wnt target genes can be expressed or inhibited and lastly the value of probe sets can be high or low.

In a BN, the (conditional) probability value or parameters for the different nodes indicate the initial belief regarding the uncertainty of an event (in some cases given another event). These parameters can be fixed manually based on user’s choice or expert knowledge or estimated using experimental data. In the reference BN, the parameter values for *β*-catenin, *TCF*4, *TRCMPLX* and the selected Wnt target genes have been set based on expert knowledge. Lastly, the parameter values for the probe sets are computed based on median thresholding of the available probe values from the training samples and addition of pseudo counts.

In this paper, training of Ref BN is done on expression values for Wnt target genes from doxycycline treated colon cancer cell line LS174T Mokry *et al*.^10^. This is done in order to have robust training of the BN which further lead to better classification. Later on, results on test data from network trained on normal and adenomas samples are compared with those obtained from Ref BN. Twelve samples were retrieved, six of which indicated Wnt signal was off and the rest showed Wnt signal was on. In six samples, were the Wnt signal was switched off, doxycycline was used to knockdown *β*-catenin in three of them. The other half that had the Wnt signal turned off was obtained using doxycycline to knockdown *TCF*4. Corresponding controls that had Wnt signal as on, were also generated when doxycycline was not used to treat either *β*-catenin or *TCF*4. For each of the twelve samples, 74 different probes were measured that gave expression values for a set of 33 different Wnt target genes. For each probe, parameter or the conditional probability values were estimated using frequency counts and median thresholding. *Thus the probability that a probe value is low given that a particular gene is off is the ratio of the number of expression values of samples treated with doxycycline that are smaller than the median of expression values of all samples*. Similarly, the probability that a probe value is high given that a particular gene is on is the ratio of the number of expression values of control samples that are greater than the median of expression values of all samples. Corresponding complements of these probabilities can be computed easily. In some cases, the probability values turn out to be zero, leading to loss of uncertainty in the interpretation. This implies that there is 100% surety that either the probe is high or low, just by measuring six samples each of Wnt on and Wnt off. *To avoid this, a pseudo count of one is added to each of the ratios in the numerator and two to the denominator, for all conditions* (gene off or on) *and states of a probe* (low or high). An example of this computation is shown in figure 4. Most of the genes get expressed when the *TRCMPLX* is active, except for a few that get inhibited. While assigning parameter values for genes during the construction of the Ref BN, certain values are reversed for genes which are inhibited when the *TRCMPLX* is active. The probability tables for the respective probesets measuring the gene expression values do not flip. This is because, the probeset values by themselves only state whether the gene is expressed or not. They do not signify whether a particular gene is inhibited or not. In order to capture the inhibited state of a gene (if it is behaving in that manner on *TRCMPLX*’s activation) its parameter values need to be flipped. While generating the inferencing engine for the Ref BN, the estimated parameter values for genes and probes (for both expressed and inhibited genes along wi) may be found swapped. This happens because, the Ref BN is unaware of the meaning of the discrete values (1 as on and 2 as off) and swapping happens randomly.

**Fig. 4.**
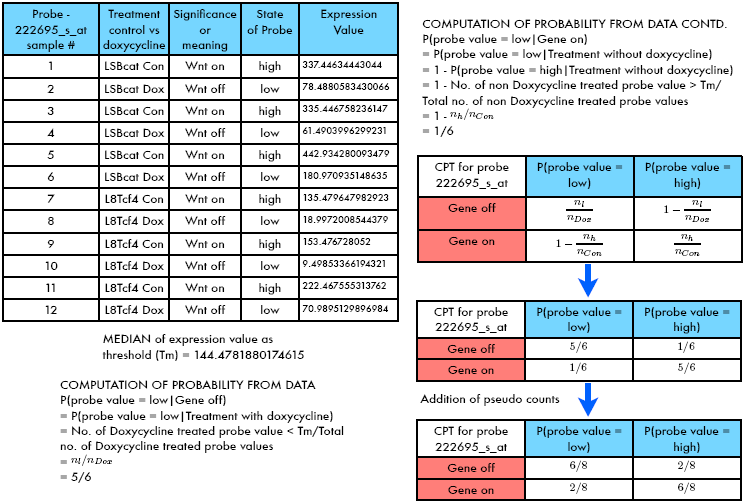
Estimation of cpd for probeset 222695-s-at for Axin2 gene, via thresholding and addition of pseudo counts.

These parameter values represent the state of gene being off or on given the *TRCMPLX* is active or inactive. This can be seen from the probability values assigned to Axin2 in figure 3. The probability values of the different nodes have been tabulated in 1. Once the state of the nodes, the causal arcs between them and the (conditional) probability tables have been defined for the Ref BN, inference need to be done. Before the inference, the inferencing engine is generated via the junction tree engine Lauritzen and Spiegelhalter^22^. The junction tree engine is used here as it forms the state of the art algorithm for exact inferencing. The current work employs the BNT toolbox Murphy *et al*. ^23^. After the engine has been generated, predictions are made for each sample in the testing data. These predictions are estimated via inferencing the state of Wnt signal being on or off. This state in turn is measured by estimating the probability of *TRCMPLX* being active, given the expression values of the probe sets of a test sample as evidence. Mathematically, the estimation is depicted in the following equation 𝓟(*TRCMPLX* = active|∀*j* probe instances *p*_*j*_ for a particular sample). A positive value of the log odds ratio of this conditional probability indicates that the Wnt signal is on and the sample is labeled as cancerous.

The predicted labels are compared with that of the ground truth labels available with the gene expression datasets to match the quality of prediction. *A similar experiment is repeated by training on normal colon and colon adenomas* (GSE8671 Sabates-Bellver *et al*.^17^) *and testing on rest of the data sets in* *table* 2. These datasets were retrieved from GSE Omnibus Edgar *et al*.^24^ and Barrett *et al*.^25^. A table similar to that shown in table 1, showing the cpd values for the network trainined on normal colon and colon adenomas is shown in tab 3. The gene expression profiles of many of the datasets were obtained from the Gene Expression Omnibus. Affymetrix GeneChip HG-U133plus2 was used for the measurement of the gene expression levels.

**Table 2.**
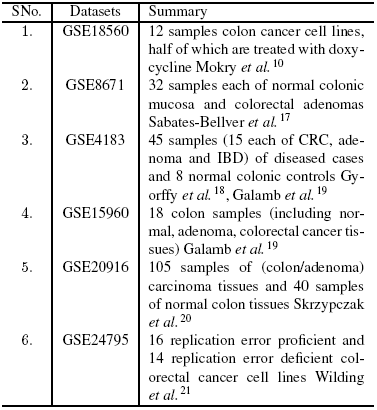
Summary of datasets used in the experiments.

| SNo. | Datasets | Summary |
| --- | --- | --- |
| 1. | GSE18560 | 12 samples colon cancer cell lines, half of which are treated with doxycycline Mokry *et al.*<sup>10</sup> |
| 2. | GSE8671 | 32 samples each of normal colonic mucosa and colorectal adenomas Sabates-Bellver *et al.*<sup>17</sup> |
| 3. | GSE4183 | 45 samples (15 each of CRC, adenoma and IBD) of diseased cases and 8 normal colonic controls Gyroffy *et al.*<sup>18</sup>, Galamb *et al.*<sup>19</sup> |
| 4. | GSE15960 | 18 colon samples (including normal, adenoma, colorectal cancer tissues) Galamb *et al.*<sup>19</sup> |
| 5. | GSE20916 | 105 samples of (colon/adenoma) carcinoma tissues and 40 samples of normal colon tissues Skrzypczak *et al.*<sup>20</sup> |
| 6. | GSE24795 | 16 replication error proficient and 14 replication error deficient colorectal cancer cell lines Wilding *et al.*<sup>21</sup> |

**Table 3.** Manually fed and estimated parameter values (cpd) for nodes of a BN constructed from normal colon and colon adenomas samples from GSE8671 Sabates-Bellver *et al*.^17^. Nodes and their parents represent the various protien complexes, genes and the corresponding probes that measure the gene expression, at different levels of the network. For normal colon and colon adenomas, genes *SLC*1*A*2, *TCF*7*L*2, *COL*18*A*1, *KLF*6 and *OAT* coloured in red indicatethat they are inhibited when *TRCMPLX* is active. Note that gene *TCF*7*L*2 measured via its corresponding probesets are not included in the colon cancer cell lines training data.

| Parameter values for nodes of Bayesian net partly using Normal colon and colon adenomas |  |  |  |  |  |  |
| --- | --- | --- | --- | --- | --- | --- |
| No. | NODES (Parents) | Reference BN (C)PTs | NODES (Parents) | Reference BN (C)PTs | NODES (parents) | Reference Bnet (C)PTs |
| 1 | B-cat | [0.40 0.60] | 222695_s.at (AXIN2) | [0.882 0.118 0.118 0.882] | 216037_x.at (TCF7L2) | [0.647 0.353 0.353 0.647] |
| 2 | TCF4 | [0.10 0.90] | 222696.at (AXIN2) | [0.971 0.029 0.029 0.971] | 216511_s.at (TCF7L2) | [0.618 0.382 0.382 0.618] |
| 3 | TRCMPLX<br>(B-CAT, TCF4) | [0.99 0.99 0.99 0.01<br>0.01 0.01 0.01 0.99] | 224176_s.at (AXIN2) | [0.971 0.029 0.029 0.971] | 210948_s.at (LEF1) | [0.589 0.411 0.411 0.589] |
| 4 | AXIN2 (TRCMPLX) | [0.95 0.30 0.05 0.70] | 224498_x.at (AXIN2) | [0.941 0.059 0.059 0.941] | 221557_s.at (LEF1) | [0.735 0.265 0.265 0.735] |
| 5 | EPHB2 (TRCMPLX) | [0.95 0.30 0.05 0.70] | 209588.at (EPHB2) | [0.912 0.088 0.088 0.912] | 221558_s.at (LEF1) | [0.735 0.265 0.265 0.735] |
| 6 | EPHB3 (TRCMPLX) | [0.95 0.30 0.05 0.70] | 209589_s.at (EPHB2) | [0.912 0.088 0.088 0.912] | 210515.at (HNF1A) | [0.50 0.50 0.50 0.50] |
| 7 | MYC (TRCMPLX) | [0.95 0.30 0.05 0.70] | 210651_s.at (EPHB2) | [0.882 0.118 0.118 0.882] | 216930.at (HNF1A) | [0.50 0.50 0.50 0.50] |
| 8 | CCND1 (TRCMPLX) | [0.95 0.30 0.05 0.70] | 211165_x.at (EPHB2) | [0.882 0.118 0.118 0.882] | 201579.at (FAT1) | [0.735 0.265 0.265 0.735] |
| 9 | SP5 (TRCMPLX) | [0.95 0.30 0.05 0.70] | 1438.at (EPHB3) | [0.882 0.118 0.118 0.882] | 226360.at (ZNR3) | [0.912 0.088 0.088 0.912] |
| 10 | LGR5 (TRCMPLX) | [0.95 0.30 0.05 0.70] | 204600.at (EPHB3) | [0.853 0.147 0.147 0.853] | 1554685_a.at (KIAA1199) | [0.912 0.088 0.088 0.912] |
| 11 | ASCL2 (TRCMPLX) | [0.95 0.30 0.05 0.70] | 202431_s.at (MYC) | [0.971 0.029 0.029 0.971] | 212942_s.at (KIAA1199) | [0.971 0.029 0.029 0.971] |
| 12 | PPARG (TRCMPLX) | [0.95 0.30 0.05 0.70] | 208711_s.at (CCND1) | [0.824 0.176 0.176 0.824] | 218704.at (RNF43) | [0.971 0.029 0.029 0.971] |
| 13 | CD44 (TRCMPLX) | [0.95 0.30 0.05 0.70] | 208712.at (CCND1) | [0.824 0.176 0.176 0.824] | 209081_s.at (COL18A1) | [0.647 0.353 0.353 0.647] |
| 14 | SLC1A2 (TRCMPLX) | [0.45 0.95 0.55 0.05] | 235845.at (SP5) | [0.912 0.088 0.088 0.912] | 209082_s.at (COL18A1) | [0.676 0.324 0.324 0.676] |
| 15 | BMP7 (TRCMPLX) | [0.95 0.30 0.05 0.70] | 210393.at (LGR5) | [0.882 0.118 0.118 0.882] | 1555832_s.at (KLF6) | [0.853 0.147 0.147 0.853] |
| 16 | TCF7L2 (TRCMPLX) | [0.45 0.95 0.55 0.05] | 213880.at (LGR5) | [0.882 0.118 0.118 0.882] | 208960_s.at (KLF6) | [0.853 0.147 0.147 0.853] |
| 17 | LEF1 (TRCMPLX) | [0.95 0.30 0.05 0.70] | 207607.at (ASCL2) | [0.882 0.118 0.118 0.882] | 208961_s.at (KLF6) | [0.794 0.206 0.206 0.794] |
| 18 | HNF1A (TRCMPLX) | [0.95 0.30 0.05 0.70] | 229215.at (ASCL2) | [0.941 0.059 0.059 0.941] | 211610.at (KLF6) | [0.647 0.353 0.353 0.647] |
| 19 | FAT1 (TRCMPLX) | [0.95 0.30 0.05 0.70] | 208510_s.at (PPARG) | [0.618 0.382 0.382 0.618] | 224606.at (KLF6) | [0.912 0.088 0.088 0.912] |
| 20 | ZNR3 (TRCMPLX) | [0.95 0.30 0.05 0.70] | 1557905_s.at (CD44) | [0.971 0.029 0.029 0.971] | 206128.at (ADRA2C) | [0.618 0.382 0.382 0.618] |
| 21 | KIAA1199 (TRCMPLX) | [0.95 0.30 0.05 0.70] | 204489_s.at (CD44) | [0.971 0.029 0.029 0.971] | 203705_s.at (FZD7) | [0.765 0.235 0.235 0.765] |
| 22 | RNF43 (TRCMPLX) | [0.95 0.30 0.05 0.70] | 204490_s.at (CD44) | [0.971 0.029 0.029 0.971] | 203706_a.at (FZD7) | [0.676 0.324 0.324 0.676] |
| 23 | COL18A1 (TRCMPLX) | [0.45 0.95 0.55 0.05] | 209835_x.at (CD44) | [0.971 0.029 0.029 0.971] | 202859_x.at (IL8) | [0.941 0.059 0.059 0.941] |
| 24 | KLF6 (TRCMPLX) | [0.45 0.95 0.55 0.05] | 210916_s.at (CD44) | [0.971 0.029 0.029 0.971] | 211506_s.at (IL8) | [0.882 0.118 0.118 0.882] |
| 25 | ADRA2C (TRCMPLX) | [0.95 0.30 0.05 0.70] | 212014_x.at (CD44) | [0.971 0.029 0.029 0.971] | 219682_s.at (TBX3) | [0.941 0.059 0.059 0.941] |
| 26 | FZD7 (TRCMPLX) | [0.95 0.30 0.05 0.70] | 212063.at (CD44) | [0.971 0.029 0.029 0.971] | 222917_s.at (TBX3) | [0.794 0.206 0.206 0.794] |
| 27 | IL8 (TRCMPLX) | [0.95 0.30 0.05 0.70] | 217523.at (CD44) | [0.853 0.147 0.147 0.853] | 225544.at (TBX3) | [0.971 0.029 0.029 0.971] |
| 28 | TBX3 (TRCMPLX) | [0.95 0.30 0.05 0.70] | 229221.at (CD44) | [0.941 0.059 0.059 0.941] | 229576_s.at (TBX3) | [0.941 0.059 0.059 0.941] |
| 29 | NKD1 (TRCMPLX) | [0.95 0.30 0.05 0.70] | 1558009.at (SLC1A2) | [0.559 0.441 0.441 0.559] | 1553115.at (NKD1) | [0.735 0.265 0.265 0.735] |
| 30 | DKK1 (TRCMPLX) | [0.95 0.30 0.05 0.70] | 1558010_s.at (SLC1A2) | [0.559 0.441 0.441 0.559] | 204602.at (DKK1) | [0.529 0.471 0.471 0.529] |
| 31 | DEFA6 (TRCMPLX) | [0.95 0.30 0.05 0.70] | 208389_s.at (SLC1A2) | [0.529 0.471 0.471 0.529] | 207814.at (DEFA6) | [0.794 0.206 0.206 0.794] |
| 32 | GLUL (TRCMPLX) | [0.95 0.30 0.05 0.70] | 225491.at (SLC1A2) | [0.588 0.411 0.411 0.588] | 200648_s.at (GLUL) | [0.706 0.294 0.294 0.706] |
| 33 | OAT (TRCMPLX) | [0.45 0.95 0.55 0.05] | 209590.at (BMP7) | [0.647 0.353 0.353 0.647] | 215001_s.at (GLUL) | [0.735 0.265 0.265 0.735] |
| 34 | LECT2 (TRCMPLX) | [0.95 0.30 0.05 0.70] | 209591_s.at (BMP7) | [0.559 0.441 0.441 0.559] | 242281.at (GLUL) | [0.441 0.559 0.559 0.441] |
| 35 | REG1B (TRCMPLX) | [0.95 0.30 0.05 0.70] | 211259_s.at (BMP7) | [0.529 0.471 0.471 0.529] | 201599.at (OAT) | [0.794 0.206 0.206 0.794] |
| 36 | SOX9 (TRCMPLX) | [0.95 0.30 0.05 0.70] | 211260.at (BMP7) | [0.471 0.529 0.529 0.471] | 207409.at (LECT2) | [0.50 0.50 0.50 0.50] |
| 37 | TDGF1 (TRCMPLX) | [0.95 0.30 0.05 0.70] | 212759_s.at (TCF7L2) | [0.706 0.294 0.294 0.706] | 205886.at (REG1B) | [0.676 0.324 0.324 0.676] |
| 38 |  |  | 212761.at (TCF7L2) | [0.706 0.294 0.294 0.706] | 202935_s.at (SOX9) | [0.824 0.176 0.176 0.824] |
| 39 |  |  | 212762_s.at (TCF7L2) | [0.559 0.441 0.441 0.559] | 202936_s.at (SOX9) | [0.971 0.029 0.029 0.971] |
| 40 |  |  | 216035_x.at (TCF7L2) | [0.618 0.382 0.382 0.618] | 206286_s.at (TDGF1) | [0.941 0.059 0.059 0.941] |

## 3 Experiments

Given the model design in figure 3, the proposed work investigates the different conditions in which the observations of the data are missing. The reason for conducting such experiments is to verify the efficacy of BNs in checking the convergence of parameters under simulated conditions and also the reproducibility of the predictions for the test data under these conditions. It should be noted that in the two runs of the training phase, one with (1) 12 samples of cancer colon cell lines (6 of which contain Wnt being on and 6 in which the Wnt is knocked down using doxycycline) (GSE18560) Mokry *et al*.^10^ and the other (2) 32 samples, each of normal colonic mucosa and colorectal adenomas (GSE8671) Sabates-Bellver *et al*.^17^, some genes are inhibited when the *TRCMPLX* is active. For the case, when 12 colon cancer cell lines are employed for training, genes *PPARG* and *KLF*6 are found to be inhibited. In the case, when normal colon and colon adenomas samples are employed, genes *SCL*1*A*2, *TCF*7*L*2, *COL*18*A*1, *KLF*6 and *OAT* are found to be inhibited. Also, apart from the set of genes used in colon cancer cell lines, gene *TCF*7*L*2, and its corresponding probeset involving 212759-s-at, 212761-at, 212762-s-at, 216035-x-at, 216037-x-at and 216511-s-at are used in the normal colon and colon adenomas training data.

In order to check the reproducibility of the parameters and the results, 5 different setups were designed. Each setup represents missing observations at different locations in the BN. Thus each setup represents a BN with a particular setting where a particular set of nodes have missing observations. A schematic representation of the setup is shown in figure 5. Training was done on each of the 5 setups based on simulated observations, sampled using the reference BN. Training was also done on each of the 5 setups based on observations obtained from real data. The reference BN was built using the real data.

**Fig. 5.**
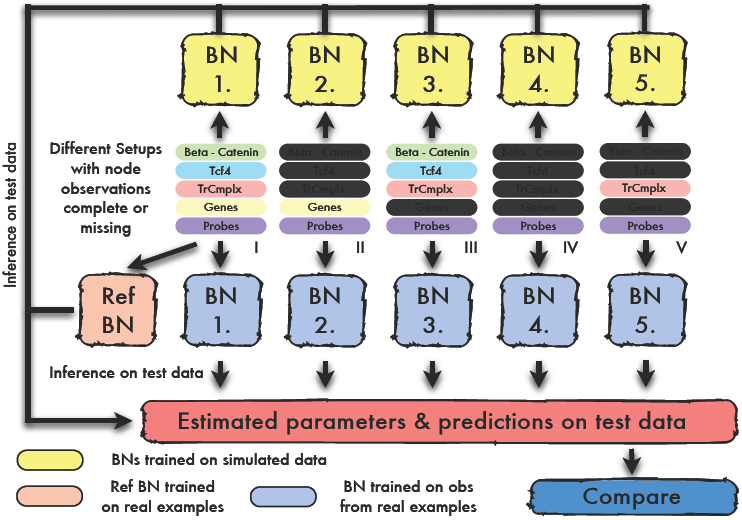
Ref BN used to infer and predict labels on test data sets. These labels matched with the ground truth labels available with the test data. Also the Ref BN is used to generate samples for all the nodes, based on the parameter values assigned to the Ref BN. In setup 1. the complete data set is present 2. observations for *β*-catenin, *TCF*4 and *TRCMPLX* are missing, 3. observations for all genes are missing, 4. except for probes, all observations for other nodes are missing and 5. observations for *β*-catenin, TCF4 and genes are missing. Missing data is shown with black coloured slabs. Parameters estimated from simulated observations and the predictions made using these parameters are compared with those obtained using the real observations for each setup. Also, parameters estimated from real observations and predictiosn made using the real data via reference BN are compared.

For simulated observations, in each setup, 1 run of training and testing was done. Training phase generated single parameter values for each of the nodes. These were then used to generate a separate prediction results for each sample in the test data. For simulated data sets, in each setup, 100 runs of training and testing were done. This generated 100 parameter values for each of the nodes in the training phase. These were then used to generate 100 separate prediction results for each sample in the test data. 95% confidence bounds were estimated on the prediction results.

The (estimated mean of the) prediction results on simulated data was compared with the prediction results obtained using the real observations for a particular setup as well as with the prediction results obtained using the reference BN, respectively. This comparison is based on checking the statistical significance of the estimated results with respect to those attained from real observations for a particular setup as well as the reference BN. (The estimated parameter values were numerically compared to see how much the deviate from those assigned to the reference BN).

The statistical significance is checked via the McNemar’s Test McNemar^26^. In a 2×2 contingency table, the McNemar’s test gives a statistic similar to the chi squared statistic which is formulated as:

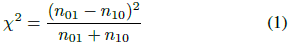

where, *n*_01_ and *n*_10_ are the false positives and the false negatives, respectively. In this study a *χ*^2^ value of 3.84 and above was set as a standard to account for the significant difference of one algorithm against another. This means that with a significance level of *α* = 0.05 and a critical value of 3.84 or less impies *p* < 0.95. For test statistic greater than the critical value 3.84, the null hypothesis is rejected, i.e. it is highly unlikely, that a setup is insignificant from the one that it is being compared with. Or the reverse, that with *p* ≥ 0.95 and a test statistic greater than 3.84, the setup is highly significant from the one it is being compared with.

### 3.1 Setup 1

As discussed in detail in section 2, setup 1 uses the Ref BN framework to train on both real and simulated observations. In this network, all nodes are observed. When using the real data, the 12 observations are derived for each of the nodes, in the network. This means that each node has 6 on/active/expressed state and 6 off/inactive/not-expressed state. For probesets, the discretized observations are made using the computed median thresholds. For a particular gene, a discretized state is determined based on majority voting on on/off states of the probes that measure its expression. The *TRCMPLX*, is assigned discretized states based on the available information that it is on for 6 doxycycline treated cases and off for 6 controls. The discretized cases for *β*-catenin and TCF4 are randomly assigned to half on and half off states. This is done because, the knowledge of the states of *TRCMPLX* explains away the effects of both *β*-catenin and TCF4. The (c)pt parameters are randomly initialized for the learner before inference can take place to generate the network engine.

In case of simulated observations, for 1 run, after the Ref BN engine has been generated the Ref BN is sampled 10000 times to generate random observations for the nodes. The (c)pt values are randomly initialized in case of a simulation run. Next, the (c)pts are learnt via inference through the maximum likelihood (ML) estimates for a simulation based new BN. The engine is generated based on the ML estimated parameters. For both the BNs (one generated based on real observations and another on simulated observations), inference is done using all probes from the testing data to predict the state of Wnt pathway.

### 3.2 Setup 2

In this setup, an exception w.r.t setup 1 is that the observations for nodes *β*-catenin, *TCF*4 and *TRCMPLX* are missing. Thus for runs on real observations, the observations are available for genes and probesets that measure the genes. The remaining experimental setup remains same as in setup 1. In case of missing observations, the (c)pts are learnt via the expectation maximization (EM) algorithm Dempster *et al*.^27^ for a new BN. Next, the engine is generated based on inference. Inference continues the same way as was done in setup 1.

### 3.3 Setup 3

The design of this setup is same as the previous setup except that the only hidden observations are for nodes for the genes. The remaining experimental setup remains same as in setup 1. In case of missing observations, the (c)pts are learnt via the expectation maximization (EM) algorithm Dempster *et al*.^27^ for a new BN. Next, the engine is generated based on inference. Inference continues the same way as was done in setup 1.

### 3.4 Setup 4

The design of this setup is same as the setup 2, except that the only observed nodes are those of the probes. An important aspect of this setup is that this is the state of the situation as it exists in reality. It implies that the only available data retrieved are the expression values. The network model built partially captures the behaviour of Wnt signaling pathway.

### 3.5 Setup 5

The design of this setup is same as the setup 3, except that the nodes *β*-catenin and *TCF*4 are also hidden.

Since the amount of training data is small (12 for colon cancer cell lines and 64 for normal colon and colon adenomas) it may happen that the estimated parameter values in the training phase may become exact (1 or 0). Thus the there would not exist a degree of belief in the happening of an event. To avoid this, an error value of 0.005 is added and the probabilities in the (c)pts are normalized to the scale of 1.

It should be noted that even though the setups are different, the dataset used is the same (leading to non independence). Thus a hypothetical test may not be valid for checking the significant differences among the various conditions imposed on a BN, trained on the same dataset. Currently, the authors are not aware of this interesting issue. In order to analyse the issue further, McNemar’s test was conducted both with the setups and between setups using predictions on different test datasets, while the training on these setups was done using two independent dataset namely (1) colon cancer cell lines GSE18560 dataset (Mokry *et al*.^10^) and (2) GSE8671 (Sabates-Bellver *et al*.^17^). Details of the analysis are presented later on.

## 4 Results

The Ref BN is trained on colon cancer cell lines GSE18560 (Mokry *et al*.^10^) with measurements of expression levels for a particular set of Wnt target genes. Perfect prediction was obtained on normal colon and colon adenoma samples from GSE8671 Sabates-Bellver *et al*.^17^. This is depicted in figure 6. In the figure, the *y*-axis depicts the log_2_ odds ratio of the probability that the *TRCMPLX* is active given the evidence that the values of the probes measured for a particular test sample (i.e. *p* = *𝓟*(*TRCMPLX*|∀*j* probe instances *p*_*j*_ for a particular sample)). The prediction in all the setups is the estimation of this marginal probability of the node *TRCMPLX* given the probe values. The log_2_ odds ratio of *p* captures the relative difference between the probability of an event occurring to the probability of the same event not occurring. Also, this relative measure marginalizes all values to a standard value of 0. Positive log odds above zero indicate probability values of the occurrence of an event being above 0.5 (i.e. 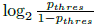 = 0 ⇒ *p*_*thres*_ = 0.5) and vice versa.

**Fig. 6.**
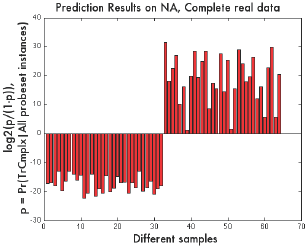
Prediction of Wnt State for 32 normal (left) and 32 adenoma (right) samples. The state of Wnt is computed via the degree of belief estimated through *p* = *𝓟r*(*TRCMPLX* is active|All probesets instances of a sample). Positive and negative values in the graph indicate adenoma and normal states, respectively. The Ref BN is trained on Colon caner cell lines.

### 4.1 Analysis of Parameter Deviation

In the simulations across the different setups, it was found that there were deviations in the estimated parameters. These deviations indicate the spread of the estimated parameters. Under different conditions depicted by the different setups, a few points can be observed. In this analysis, parameter values for two genes with their corresponding probesets have being depicted. In each of the graphs in the following figures 7 and 8, the y-axis depicts the probability of the node being 1 or off when the parent node is 2 or on and the x-axis depicts the probability of the node being 1 or off when the parent node is 1 or off. The motivation behind comparing these two probabilities is to see how much reproducible the parameter estimates are for a particular state of the node (here the gene under consideration) under the different states in which the parent node can be.

**Fig. 7.**
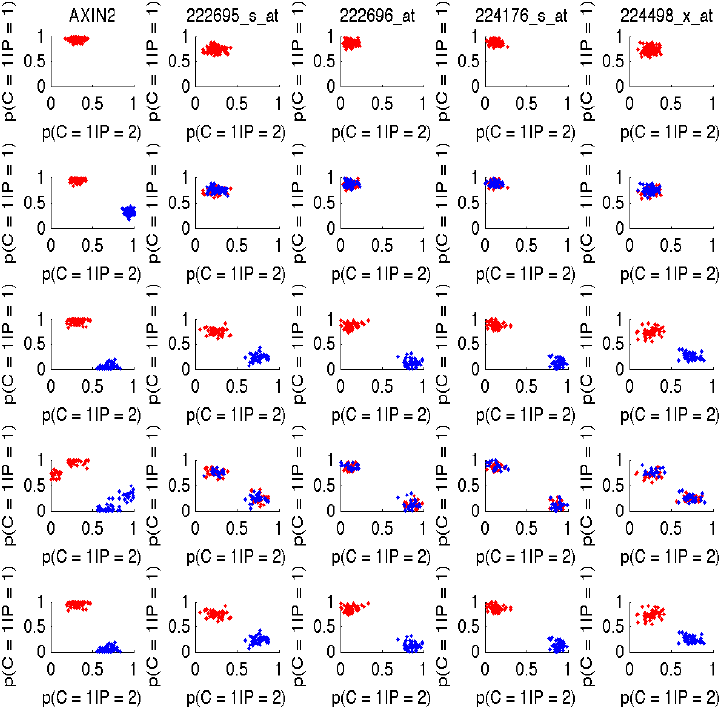
Plot of deviations in estimated parameter for AXIN2 and its corresponding probesets. Rows represent the different setups. Columns indicate the node name on the top. x-axis indicates the probability of child being off given that the parent is off i.e. p(C = 1|P = 2). y-axis indicates the probability of child being off given that the parent is on i.e. p(C = 1|P = 1).

**Fig. 8.**
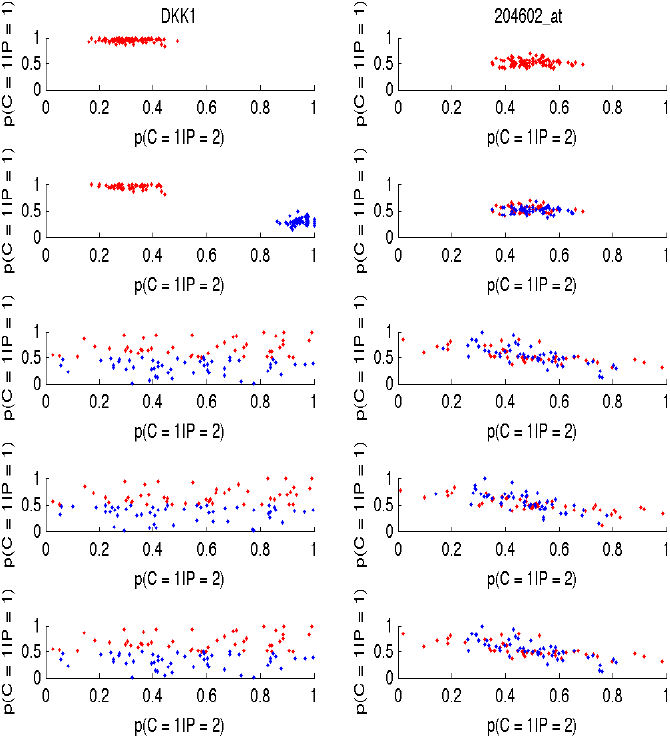
Plot of deviations in estimated parameter for DKK1 and its corresponding probesets. Rows represent the different setups. Columns indicate the node name on the top. x-axis indicates the probability of child being off given that the parent is off i.e. p(C = 1|P = 2). y-axis indicates the probability of child being off given that the parent is on i.e. p(C = 1|P = 1).

In the graphs shown in the above mentioned figures, the red coloured points indicate how close the estimated parameters are to parameter values assigned to the node under consideration, in the reference network. The blue points indicate the flip in the estimated parameter values while learning the parameter on the bayesian network. The flips occur mainly due to the fact that while estimating the parameters for parent nodes with hidden (or no parameter) values, the bayesian network is unable to decide the which value to assign to a particular state of the child node. In this study, in a hundred simulations, these flipping can happen along rows of the cpt or along the columns of the cpt. It is also expected that if the flipping occur, they will happen approximately half of time as there are only two discretized stated for a node and the bayesian network would provide solutions for both normal and flipped cases, equally.

Considering column one in figure 7, the deviations in estimated parameters for AXIN2 have been tabulated for setups one to five from top to bottom. Using setup 1, when all the nodes are observed, the estimated value of the parameters (i.e p(AXIN2 = 1 or off *TRCMPLX* = 1 or off) and p(AXIN2 = 1 or off | *TRCMPLX* = 2 or on)) spread around the values (0.95 and 0.30) of the same parameters assigned in the reference bayesian network. In the second row and first column, with the the *TRCMPLX* being hidden, flipping of the values of the parameters occur for AXIN2. This is evident in second row and first column graph of the same figure. In the third setup, the nodes for the genes is hidden. Thus the estimation of the parameters gets flipped around the column of the cpt table as this time the bayesian network does not know the distinction between whether discrete state 1 means a gene off or on (vice versa for discrete state 2). In the forth setup, both the genes and the *TRCMPLX* are hidden, thus flipping happens both at row level and column level. This is clearly evident from the two cluster formations in both the red and blue regions. Lastly, setup five is similar to setup three as knowledge of information about *TRCMPLX* explains away the effects induced by *β*-catenin and TCF4. Thus the bottom graph looks similar to the graph in the third row.

Considering graphs in a row of figure 7, the parameter values of the corresponding probesets show similar behaviour based on that shown by the respective gene for which the former encode the activation values. The mixed cluster of blue dots and red dots for the probesets point to number of parameter values for probes for which flipping of parameter values for the genes have happened. In case the probeset values have non discriminative noisy parameter values, then it is often found that estimated parameter values for the genes (if the gene node is hidden) are completely noisy. This can be seen in the graphs for setups 3, 4, and 5 in figure 8. In rows 1 and 2 for the same figure, the spread in the parameter values for the genes do not show any affect even if the parameter values for the probesets are noisy. This is due the fact that the nodes for the genes are observable. Thus these deviation plots point to the reproducibility of the parameter values. Also, for noisy probeset values, it is not possible to estimate good parameter values for their respective genes, in case the gene nodes are hidden. The deviation study shows a way of the sensitivity of the estimated parameter values for each and every node under different setups using the bayesian network. Finally, deviations across all setups for each of the nodes for one of the simulation run is depicted in figures 9 and 10.

**Fig. 9.**
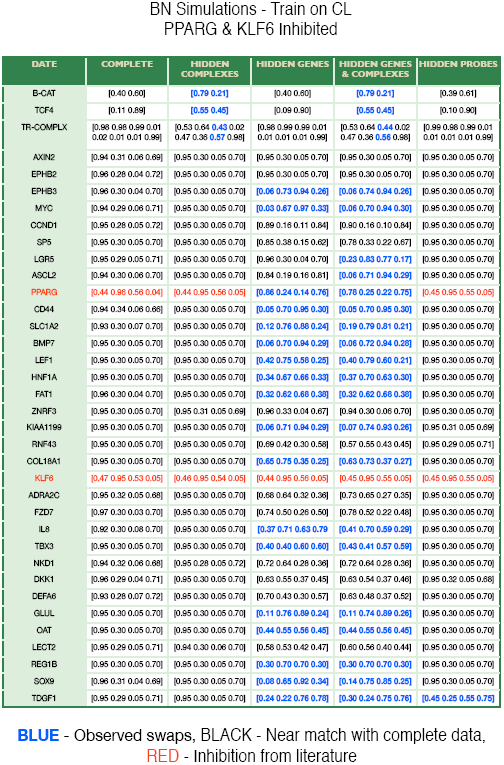
Deviation of parameters using GSE18560 dataset (Mokry *et al*.^10^)

**Fig. 10.**
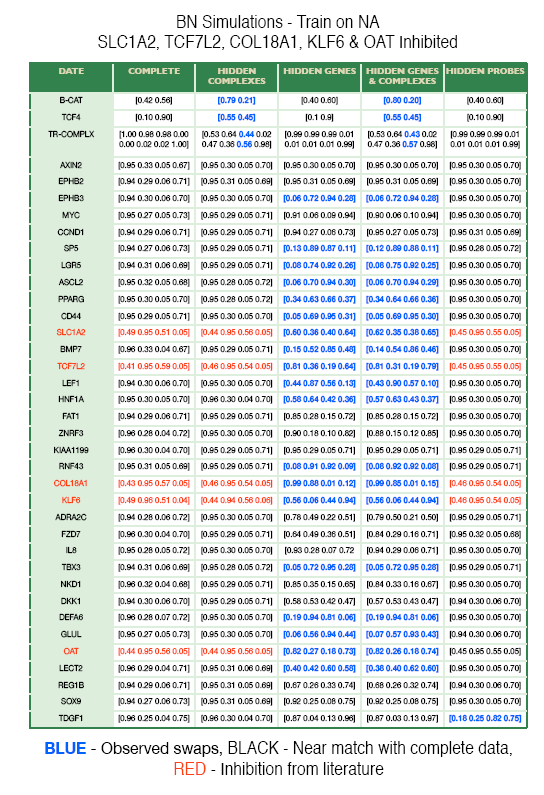
Deviation of parameters using GSE8671 (Sabates-Bellver *et al*.^17^)

### 4.2 Results from setups based on real observations

Next for each of the setups, the real data was used to generate discretized observations and depending on complete or missing observations, ML or EM algorithm was employed to learn the (c)pt values. These were then used to predict the state of samples as normal colon or colon adenoma. The predictions for each of the setup using the real data set is shown in figure 11. In the graphs for predictions made via estimates computed from observations, a pseudo count of 10^−10^ is added to the probability values in order to circumvent the problem of null behaviour (log_2_ odds computes to zero as prediction evaluates to 0.5) or infinite relative difference (log_2_ odds computes to ∞) that afflicts the graphical depiction.

**Fig. 11.**
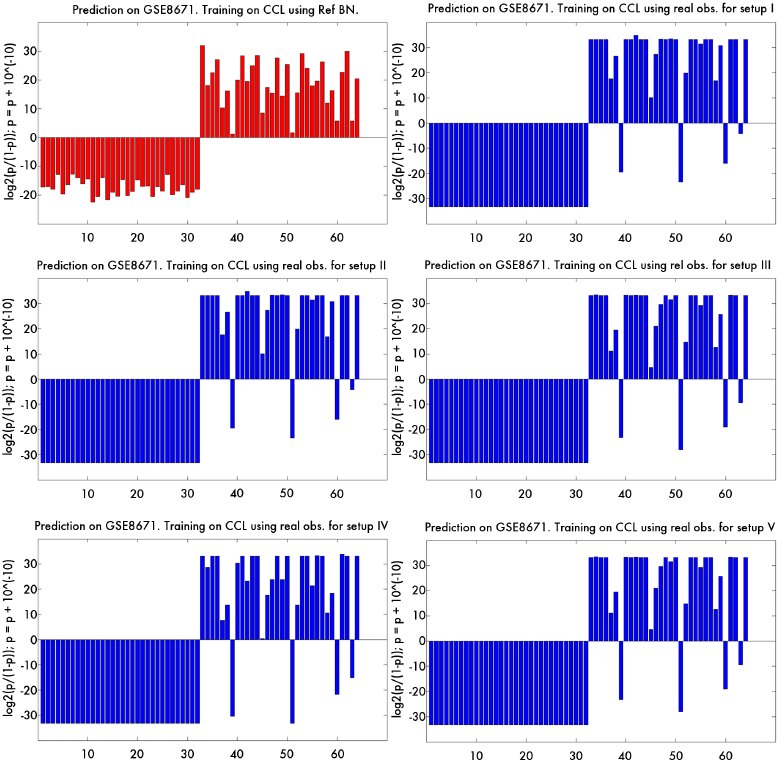
Description same as in figure 6. Left to right row wise, the graphs depict the predictions for BNs obtained from (a) complete data (Ref BN) (b) complete observations (BN from setup 1) (c) observed probes and genes (BN from setup 2) (d) observed probes and complexes (BN from setup 3) (e) observed probes (BN from setup 4) and (f) observed probes and *TRCMPLX* (BN from setup 5). BNs for all setups here use real observations.

Comparing the predictions obtained via the observations in different setups to that obtained via the predictions obtained from Ref BN using real data, shows that the results do not deviate significantly. This is depicted in the plots of figure 11. It can be seen that the EM based estimates generate misclassifications on samples (for example the 50^*th*^ sample) that are nearly classified to colon adenomas by the reference BN using complete data. The misclassifications may be attributed to the probeset values for those samples that do not satisfy the threshold criterion by a negligible amount. The predictions may appear to be same due to the clipping effect induced by addition of pseudo counts to the probability values, but in reality are not the same. Thus, non significance does not imply equality in prediction. Even though the label assigned in most cases may be same, the relative values of the estimated probabilities and their respective log odds can be different (example, the log odds ratio of the prediction on 50^th^ sample in graphs for all setups generated from real observations show dissimilar values but same assigned label).

This non significance was indicated by the *χ*^2^ value generated via the McNemar’s test using equation 1, for all predictions obtained through each of the BNs from all the setups vs the predictions obtained via the Ref BN. The respective *χ*^2^ values per setup have been depicted in column 2 of table 4. All values lying below 3.84, indicate that the probability of the deviation being high is lower that 0.95. Thus with an *β* = 1.05, the null hypothesis holds i.e. the deviations of predictions from BNs representing different setups based on real observations are insignificant from predictions made using Ref BN. It must be noted that the insignificance does not imply equality in results.

**Table 4.** Comparison 1. BNs based on Obs. from real data VS Ref BN based on real data (a between setup comparison). Comparison 2. BNs based on Simulated Obs. VS Ref BN based on real data (a between setup comparison). Comparison 3. BNs based on Simulated Obs. from real data VS BNs based on Obs. from real data (a within setup comparison). McNemar’s Test with significance level *α* = 0.05 and *χ*^2^ value of 3.84 ⇒ *p* < 0.95 with 1 degree of freedom. Values higher than the critical value imply significant differences with a *p* ≥ 0.95.

| Comp. 1 | Test | Comp. 2 | Test | Comp. 3 | Test |
| --- | --- | --- | --- | --- | --- |
| Setup I | 2.25 | Setup I | 0 | Setup I | 2.25 |
| Setup II | 2.25 | Setup II | 0 | Setup II | 2.25 |
| Setup III | 2.25 | Setup III | 0.5 | Setup III | 0.5 |
| Setup IV | 2.25 | Setup IV | 0.5 | Setup IV | 0.5 |
| Setup V | 2.25 | Setup V | 0.5 | Setup V | 0.5 |

Finally, table 5 shows the deviations in predictions using the real observations with respect to predictions obtained using the Ref BN. The error is computed using the norm-2 formulation, i.e. if 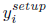 is the prediction generated using a particular setup and 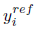 is the prediction generated using the ref BN for *i*^*th*^ sample:

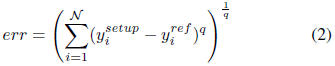

were *q* = 2. Besides the err, the avg err which is the mean of the err in equation 2 and the std err which is the standard deviation of the norm-1 (*q* = 1) err is also computed. The average error is depicted in %. It can be seen that most predictions made using the real observations has been clipped by addition of pseudo count to the estimated *p* values. Thus most of the estimated error depicted in the table show similar values, which may not be the case. These error do not point to missclassification but only to the value of prediction that is being made using the real observations with respect to the prediction made using the reference network. The major deviations may be accounted for the fact that the BN trained on observations contain very less amount of training data which may not lead to good parameter estimation during the training phase. Due to not so good parameter estimates, it is possible that the prediction values on sample even though correct may have large deviations from the prediction values obtained using Ref BN. This is supplemented by the fact that, error computed using predictions from simulation are much lower. This can be seen in the tabulated values of the err, avg err and std err from predictions using simulation in the same table. A reason for close predictions via simulation is due to the fact that observations are sampled from the same ref BN. But the error deviations jump by two to three times when using setups III, IV and V, in comparison to deviations obtained from setups I and II. Details of peculiar behavior in deviations generated using simulations are discussed later.

**Table 5.** Error measurements generated from deviation of predictions made via BNs based on simulations and real observations for different setups with respect to the predictions made via the ref BN. Here err denotes the norm-2 error, avg err denotes the average of the norm-2 error and std err denotes the standard deviation of the error per sample computed using norm-1.

| Predictions of Sim. BN vs Ref BN |  |  |  |
| --- | --- | --- | --- |
| Setups | err | avg err in % | std of err |
| I | 6.4862 | 10.13 | 0.6802 |
| II | 6.7011 | 10.47 | 0.5767 |
| III | 20.2022 | 31.56 | 1.6364 |
| IV | 302.08 | 4.7200 | 12.0053 |
| on inversion | 15.4472 | 0.2413 | 0.9253 |
| V | 20.2023 | 0.3156 | 1.6364 |
| Predictions of Real obs. BN vs Ref BN |  |  |  |
| Setups | err | avg err in % | std of err |
| I | 112.93 | 176.46 | 5.3649 |
| II | 112.93 | 176.46 | 5.3643 |
| III | 113.09 | 176.71 | 6.2144 |
| IV | 114.09 | 178.26 | 7.1350 |
| V | 113.09 | 176.71 | 6.2140 |

### 4.3 Results from setups based on simulated observations

Once the predictions have been made using real observations, another run across the setups is done using observations sampled from respective BNs through simulation. The interpretations of the graphs and the McNemar’s test remain the same as in the previous subsection. Similar significance results were obtained while comparing the predictions from BNs based on sampled observations from simulation in respective setups apropos to predictions from Ref BNs. These significance values can be viewed in the 4^th^ column of table 4. Figure 12 depict the prediction results obtained using BNs based on the sampled observations for each of the setups. It should be noted that the error rates computed from deviations

**Fig. 12.**
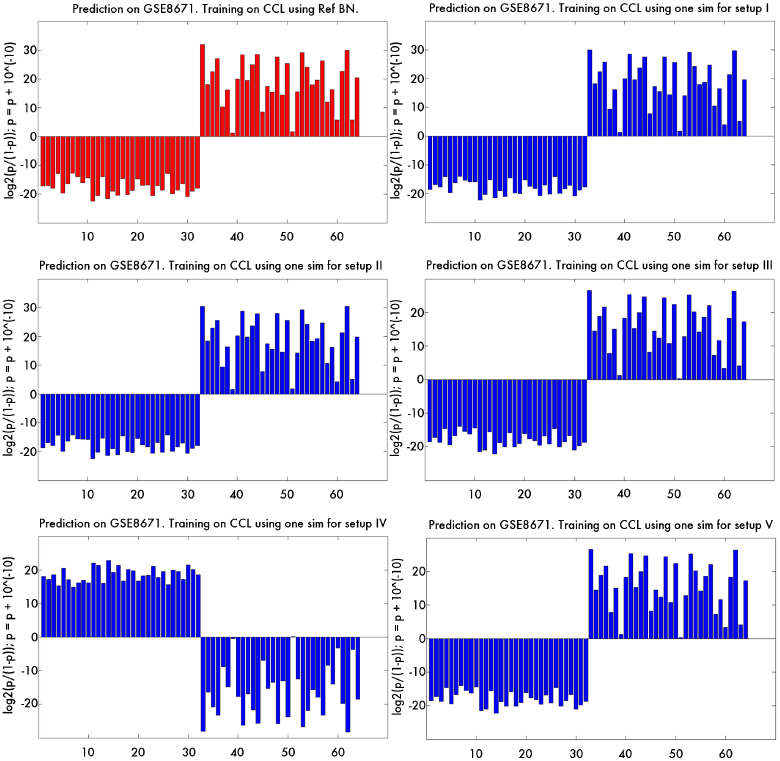
Description same as in figure 6. Left to right row wise, the graphs depict the predictions for BNs obtained from (a) complete data (Ref BN) (b) complete observations (BN from setup 1) (c) observed probes and genes (BN from setup 2) (d) observed probes and complexes (BN from setup 3) (e) observed probes (BN from setup 4) and (f) observed probes and *TRCMPLX* (BN from setup 5). BNs for all setups here use sampled observations from simulation.

Finally, 5 shows the prediction deviations computed using the BNs based on simulations with respect to predictions given by the Ref BN. One of the peculiar result is the error obtained using the setup IV, where the predictions are actually inverted as can be seen in figure 12. The flip mainly occurs due to the fact that nodes for both the *TRCMPLX* and the genes are hidden in case of simulations. In such a scenario, while estimating the parameter values, as can be seen in the forth row of figure 7 for example, there is no clear indication of a parameters converging to those assigned in the Ref BN, with flips happening both row (i.e. at *TRCMPLX* level) and column wise (i.e. at gene level). This leads to inversion in the prediction values of the samples in the test case. To see the actual error, the generated prediction values were inverted and err was recomputed. These inverted values then point to deviations that are near to those predictions made by the Ref BN. Also, hiding genes in setups III, IV and V often lead to greater deviations in predictions. This is because, if probeset values are non discriminative and noisy (as in figure 8), then the estimation of parameters for genes is also noisy. With a BN based on noisy set of parameters, it may happen that the inference on the state of *TRCMPLX* given the new test sample readings (that is the probeset values for a sample) may give prediction values, which are far from those estimated by the Ref model where the assigned values of genes are based on expert knowledge.

### 4.4 Comparing predictions from simulated vs real observations

Lastly, for each of the setups, the statistical significance of the prediction results within the setup also need to be compared. This is done to insure the quality of reproducibility based on real and simulated observations. The values found in 6^*th*^ column of table 4 indicates the significance between the results. It is found that prediction deviations generated from simulated observations are not significant with respect to those generated via the real observations.

It is imperative to note that non-significance does not imply equality in results. Prediction values can be dissimilar for a particular sample but the assigned label can be same. The McNemar’s test captures the deviation in prediction labels and not the deviations in the prediction values. Thus similarity in BN results do not imply equality. In case of using observations when the complete set of nodes is observable, the prediction labels match the ground truth values for NA. Thus the BN is found to be accurate in labeling the samples given the full set of nodes is observable. These minute aspects can be seen in the graphs of figures 11 and 12.

Finally, the deviations in the predicted probability values for the 53 samples of the test data set GSE4183 have been plotted. This is done to check the reproducibility of the prediction results for 100 simulations in each of the setups. Figure 14 shows the box plot measurements of the predicted probabilities for each of the sample of the GSE4183 dataset. Table 6 is refers to the error in predictions made while using the setups with respect to those made by the RefBN. In figure 14, the box plots represent the spread of the probability values that represent Pr(*TRCMPLX* = on | all instances of probeset values per sample). The red bar within the blue boxes represent the median while the bottom and the top portions of the boxes represent 25 and 75 percentile of the data. The whiskers beyond the boxes represent the extreme areas up to which the predictions spread. The plus symbols in red colour indicate the outliers that are not a part of the spread. It can be seen that the deviations in the predicted probability values is less for setups I, III and V. Setup V gives similar results as compared to setup III, because the observations of *TRCMPLX* makes the readings of *β*-catenin and TCF4 redundant. The predictions deviate much more for setups II and IV (figures 15 and 16). This can be attributed to the fact that when the *TRCMPLX* is hidden, then estimation of its parameters may not depict probability values close to those in the Ref BN. Using simulated BNs based on these estimated parameters leads to large deviations in predicted probabilities. These are then reflected with high average percentage error in table 6. Table 6 shows the deviations in the error over 53 samples and the average percentage error with respect to predictions made using the Ref BN. Clearly, setups I, III and V outperform setups II and IV. But a slight strange result is also noticed when setup I gives slightly bad result compared to setups III and V. We could not come up with as reasonable answer for this behaviour.

**Fig. 13.**
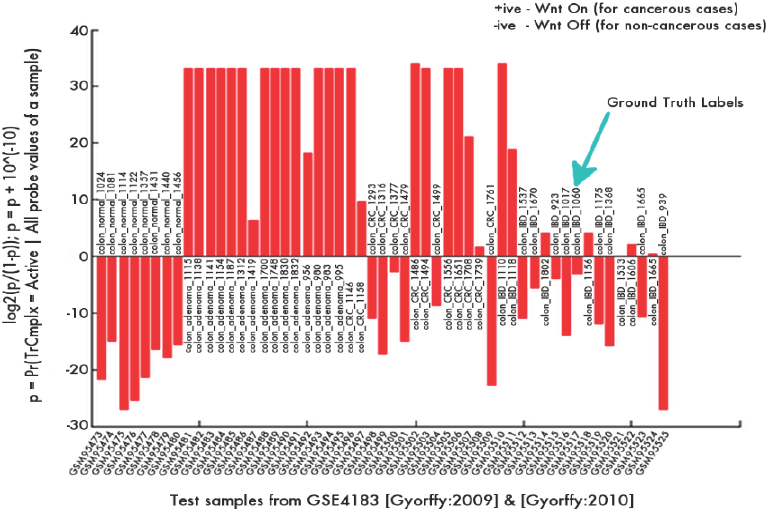
Y-axis - Log ratios of marginal probability of *TRCMPLX* being active (indicating Wnt on) on samples from GSE4183. Inference was based on Ref BN trained on 64 samples of GSE8671 (Sabates-Bellver *et al*.^17^).

**Fig. 14.**
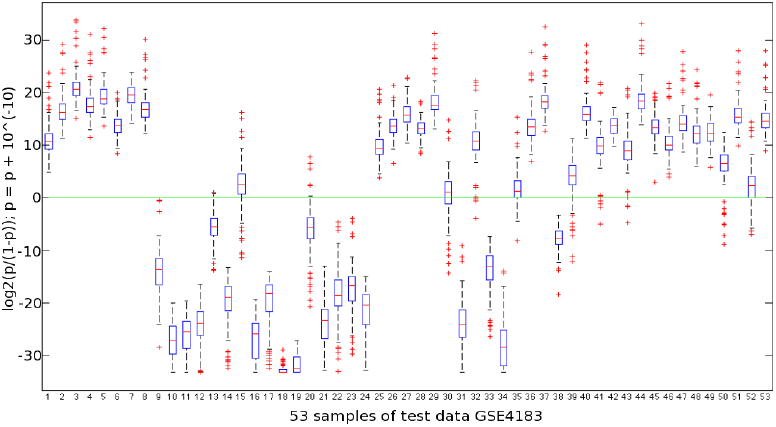
Box plot of the log2 odds of predicted p = Pr(*TRCMPLX* = on | all instances of the probesets are available), using 100 simulations in setup I for all samples in GSE4183. The blue boxes indicate the 25 and 75 percentile. The red line in the blue boxes represent the median value. The whiskers represent the extreme points and the red plus symbols represent the outliers.

**Fig. 15.**
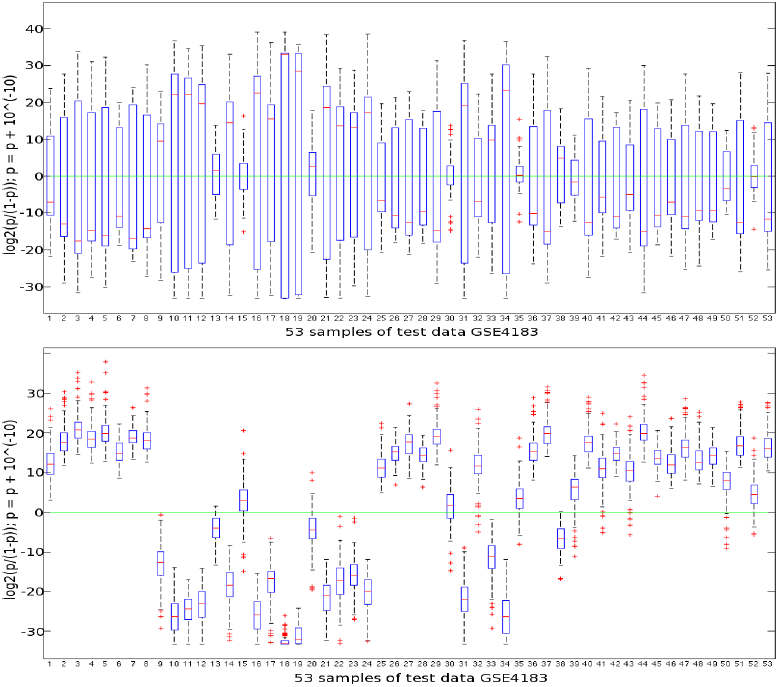
Box plot of the log2 odds of predicted p = Pr(*TRCMPLX* = on | all instances of the probesets are available), using 100 simulations in setup II (top) and setup III (bottom) for all samples in GSE4183. Descriptions remain same as in figure 14

**Fig. 16.**
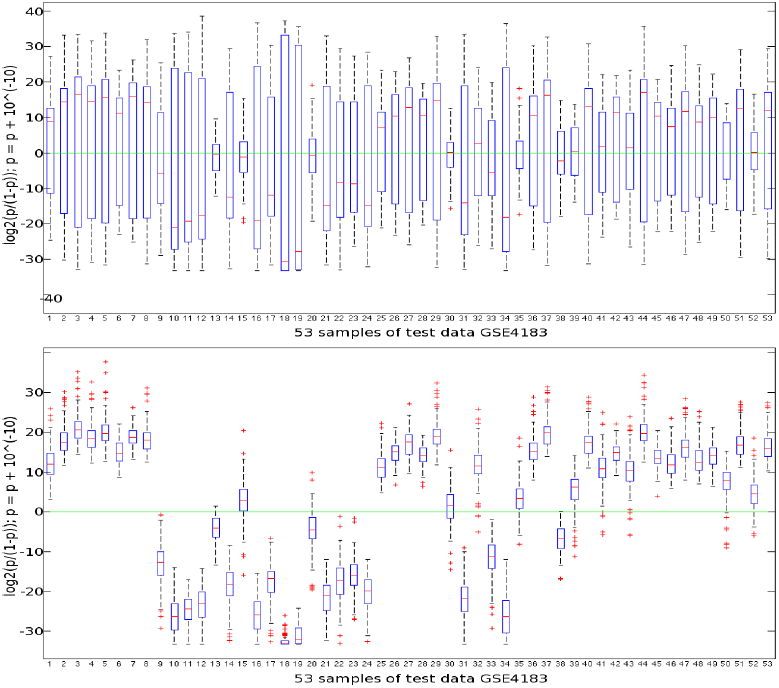
Box plot of the log2 odds of predicted p = Pr(*TRCMPLX* = on | all instances of the probesets are available), using 100 simulations in setup IV (top) and setup V (bottom) for all samples in GSE4183. Descriptions remain same as in figure 14.

**Table 6.** Average error w.r.t predictions made by the corresponding RefBN trained on colon cancer cell lines and the standard deviation in the error for each of the setups has been shown.

| Log-odds Error Statistics w.r.t Ref BN |  |  |
| --- | --- | --- |
|  | Avg. Err | Std. in Err |
| Setup I | 1.3539 | 1.9547 |
| Setup II | 328.68 | 298.87 |
| Setup III | 2.6005 | 2.6995 |
| Setup IV | 211.87 | 191.82 |
| Setup V | 2.5835 | 2.6847 |

### 4.5 Prediction results on various test data sets

Next, the focus is shifted to evaluating the quality of results for samples obtained from different types of cancer. The list of these datasets are tabulated in table 2. The prediction results for Ref BN trained on two different datasets and tested on the tabulated data in table 2 are show in table 8. In table 8, the tabular results on top are predictions from Ref BN trained on 12 (6 each from doxycycline treated Wnt pathway i.e. Wnt off and controls i.e. Wnt on) samples from Colon cancer cell lines GSE18560 (Mokry *et al*.^10^). The bottom tabular results depict predictions from Ref BN trained on G4 (32 each of normal colon and colon adenomas) samples from GSE8671 Sabates-Bellver *et al*. ^17^.

**Table 7.** Contingency table for a binary problem

|  |  | Ground Truth |  |
| --- | --- | --- | --- |
|  |  | + | - |
| Predicted<br>label | + | $TP$ | $FP$ |
| | - | $FN$ | $TN$ |

**Table 8.**
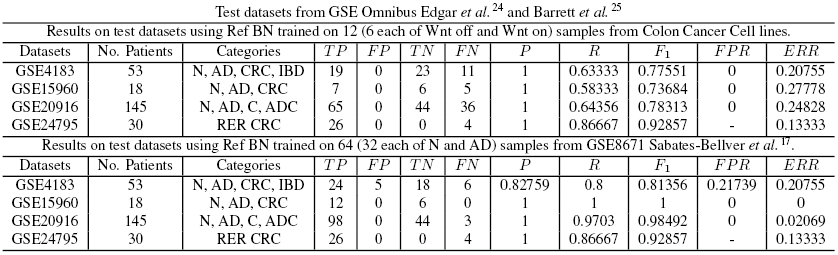
Abbreviations : N - normal, AD - adenomas, CRC - colorectal cancer, IBD - inflammatory bowel diseases, C - carcinomas, ADC - adenocarcinomas, RER CRC - replication error colorectal cancer.

Before, discussing the details of the table and the evaluated measures, a succinct analysis of one of the prediction results on 53 samples from GSE4183 Gyorffy *et al*.^18^ and Galamb *et al*.^19^, is presented in figure 13. Among the 53 test samples, the first 8 represent normal (N) cases, the second 15 represent adenoma (AD) cases, the third 15 represent colorectal cancer (CRC) cases and last 15 represent inflamatary bowel disease (IBD) cases. These ground truth labels were retrieved from respective GSE profile from GSE Omnibus Edgar *et al*.^24^ and Barrett *et al*. ^25^. These labels for these samples are denoted in the previously mentioned figure, along with the log ratios. To reiterate, +ive log odds indicate Wnt on and vice versa. Thus, it is expected that for AD and CRC cases the log odds are +ive and for N and IBD cases, the log odds are –ive. Given the predications on this test data, the quality of the predictions need to be assessed.

This assessment is done via a list of measures, a description of which follows. Contingency matrix values were used to derive the measures. A contingency matrix compares how the predicted observations behave with respect to the observed ground truth labels. Four measures indicate the match and deviations between the predictions and the ground truth labels. These are *true positive* or *TP*, *false positive* or *FP*, *true negative* or *TN* and false negative *FN*, shown in table 7.

Next, the precision *P* of the prediction results indicate how precise the labels are and is estimated via 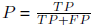. This does not mean that the results are accurate. Accuracy can be measured using the total error computed via the final error in predictions estimated via 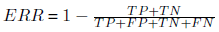. The recall *R* of the results indicate how many of the labels were correctly retrieved from the original set of true labels (*R* = 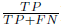). To measure the effectiveness of retrieval, an *F*_*β*_ score is evaluated that gives β times as much weightage to precision as recall. Here *β* was taken as 1. The *F*_*β*_ is computed as follows: 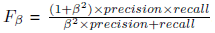. Finally the false positive rate *FPR* is also reported. The *FPR* is the probability of falsely rejecting the null hypothesis that the samples are normal cases and is computed via 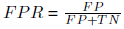. The estimated values for each of the datasets can be seen in table 8.

As can be seen from the results on GSE4183 (predictions based on Ref BN trained on GSE8671 samples), high value (nearing to 1) of *F*_1_-score (0.81356) indicates both high precision (0.82759) and high recall (0.8). High precision points to retrieval of high concentration of predictions. High recall points to retrieval of maximum number of originally relevant data. While giving equal weightage to both precision and recall, the high *F*_1_-score ensure that the quality of predictions is good. Also, the low total error 0.20755 on the test data indicate that prediction levels are nearly accurate using the Ref BN trained on GSE8671. Finally, a low value of *FPR* (0.21739) indicate the probability that less number of examples are falsely derived as positive cases of carcinoma in colon.

Also, while comparing the results obtained from Ref BN trained on 12 colon cancer cell lines GSE18560, a low *F*_1_-score with respect to that obtained from Ref BN trained on 64 normal colon and colon adenomas of GSE8671, indicate that either the low number of samples is not enough to correctly classify the test data or data sets from controlled settings (colon cancer cell lines treated with doxycycline) do not capture the actual phenomena and thus give low prediction results. The latter case holds with much more weight as the number of *FN* is higher for predictions obtained from colon cancer cell lines GSE18560 than those obtained from GSE8671.

Overall, from all the test datasets assessed, the results obtained from Ref BN trained on GSE8671 are much better that those obtained from Ref BN trained on colon cancer cell lines GSE18560. These indications are magnified in case of datasets with large number of test examples. Taking the case of GSE20916 Skrzypczak *et al*.^20^, the number of false negatives has drastically reduced from 44 (while using Ref BN trained on colon cancer cell lines GSE18560) to 3 (while using Ref BN trained on GSE8671). There is a corresponding increase in recall from 0.64356 to 0.9703 and *F*_1_-score from 0.78313 to 0.98492. It must be noted that precision value remains the same for both the results. This gives another insight into why *F*_*β*_-scores are important in accessing the quality of retrieved results. Lastly, the final estimated error reduces significantly from 0.24828 to 0.02069.

Finally, coming back to the issue of validity of conducting McNemar’s test on setup designs which are different yet trained on same data, it was found that hypothesis testing may not be the correct way to prove the statistical significance of setup designs. As expected from table 4, most of the setups were found to be statistically insignificant from each other as well as with the Ref BN, when the training set remained the same. A further analysis of within setup and between setups was also conducted. All setups where trained on two different independent datasets namely (1) colon cancer cell lines GSE18560 Mokry *et al*.^10^ and (2) GSE8671 Sabates-Bellver *et al*.^17^. The evaluated McNemar values were generated using independent test datasets mentioned previously. Table 9 shows how the significance values behave with different datasets. What can be inferred from these values is described next.

**Table 9.** CCC implies colon cancer cell. McNemar values obtained from comparing BNs for setups. Both within setup and between setup comparison is done. McNemar’s Test with significance level *α* = 0.05 and *χ*^2^ value of 3.84 ⇒ *p* < 0.95 with 1 degree of freedom. Values higher than the critical value imply significant differences with a*p* ≥ 0.95.

| Trained on<br>GSE8671 →<br>CCC lines ↓ | Test dataset GSE4183 Györfy *et al.*<sup>18</sup>, Galamb *et al.*<sup>19</sup> |  |  |  |  |  |
| --- | --- | --- | --- | --- | --- | --- |
|  | Ref<br>BN | Setup I<br>BN | Setup II<br>BN | Setup III<br>BN | Setup IV<br>BN | Setup V<br>BN |
| Ref BN | 10.000 | 0.333 | 5.000 | 0.333 | 5.000 | 0.333 |
| Setup I BN | 12.000 | 3.000 | 7.000 | 3.000 | 7.000 | 3.000 |
| Setup II BN | 12.000 | 3.000 | 7.000 | 3.000 | 7.000 | 3.000 |
| Setup III BN | 12.000 | 3.000 | 7.000 | 3.000 | 7.000 | 3.000 |
| Setup IV BN | 12.000 | 3.000 | 7.000 | 3.000 | 7.000 | 3.000 |
| Setup V BN | 12.000 | 3.000 | 7.000 | 3.000 | 7.000 | 3.000 |
| Trained on<br>GSE8671 →<br>CCC lines ↓ | Test dataset GSE15960 Galamb *et al.*<sup>19</sup> |  |  |  |  |  |
|  | Ref<br>BN | Setup I<br>BN | Setup II<br>BN | Setup III<br>BN | Setup IV<br>BN | Setup V<br>BN |
| Ref BN | 5.000 | 4.000 | 1.000 | 0.667 | 2.000 | 0.667 |
| Setup I BN | 7.000 | 1.000 | 0.000 | 0.000 | 4.000 | 0.000 |
| Setup II BN | 7.000 | 1.000 | 0.000 | 0.000 | 4.000 | 0.000 |
| Setup III BN | 7.000 | 1.000 | 0.000 | 0.000 | 4.000 | 0.000 |
| Setup IV BN | 7.000 | 1.000 | 0.000 | 0.000 | 4.000 | 0.000 |
| Setup V BN | 7.000 | 1.000 | 0.000 | 0.000 | 4.000 | 0.000 |
| Trained on<br>GSE8671 →<br>CCC lines ↓ | Test dataset GSE20916 Skrzypczak *et al.*<sup>20</sup> |  |  |  |  |  |
|  | Ref<br>BN | Setup I<br>BN | Setup II<br>BN | Setup III<br>BN | Setup IV<br>BN | Setup V<br>BN |
| Ref BN | 33.000 | 0.818 | 6.250 | 3.267 | 21.160 | 3.267 |
| Setup I BN | 36.000 | 1.500 | 6.259 | 5.000 | 24.143 | 5.000 |
| Setup II BN | 36.000 | 1.500 | 6.259 | 5.000 | 24.143 | 5.000 |
| Setup III BN | 37.000 | 2.130 | 7.538 | 5.762 | 25.138 | 5.762 |
| Setup IV BN | 37.000 | 1.960 | 7.000 | 5.762 | 27.000 | 5.762 |
| Setup V BN | 37.000 | 2.130 | 7.538 | 5.762 | 25.138 | 5.762 |
| Trained on<br>GSE8671 →<br>CCC lines ↓ | Test dataset GSE24795 Wilding *et al.*<sup>21</sup> |  |  |  |  |  |
|  | Ref<br>BN | Setup I<br>BN | Setup II<br>BN | Setup III<br>BN | Setup IV<br>BN | Setup V<br>BN |
| Ref BN | 6.000 | 5.000 | 5.000 | 5.000 | 5.000 | 5.000 |
| Setup I BN | 10.000 | 9.000 | 9.000 | 9.000 | 9.000 | 9.000 |
| Setup II BN | 10.000 | 9.000 | 9.000 | 9.000 | 9.000 | 9.000 |
| Setup III BN | 11.000 | 10.000 | 10.000 | 10.000 | 10.000 | 10.000 |
| Setup IV BN | 11.000 | 10.000 | 10.000 | 10.000 | 10.000 | 10.000 |
| Setup V BN | 11.000 | 10.000 | 10.000 | 10.000 | 10.000 | 10.000 |

From a global perspective, for each of the test dataset in table 9, it can be seen that Ref BN trained on GSE8671 is significantly different from all the BNs trained on colon cancer cell lines GSE18560 (2^*nd*^ column). This is not the case with the Ref BNs trained on colon cancer cell lines (and tested on each of the four independent datasets) which is significantly different for some of the setups and significantly not different from other setups (rows 3,11,19, 29). One may draw a conclusion that the GSE8671 datasets capture more properties than the colon cancer cell lines GSE18560 and are thus give good performance in comparison to the later.

Similar consistent behaviour (as Ref BN trained on GSE8671) can be found for Setup IV BN trained on GSE8671 (6^*th*^ column). The consistency of the results obtained from Setup IV also points to the fact that even when observations for probes are available (estimated from real dataset), it is able to point to the significant difference between the GSE8671 and colon cancer cell lines GSE18560 dataset. This fact gets blurred in the other design setups where the McNemar values are not consistent within and between setups. Values from Setup I while using GSE4183, GSE15960 and GSE20916 show that GSE8671 is not significantly independent from colon cancer cell lines GSE18560 (except in the case of GSE24795). Similarly, values from Setup II while using GSE4183, GSE20916 and GSE24795 show that GSE8671 is significantly independent from colon cancer cell lines data. For Setups III and V, using GSE4183 and GSE15960, the data sets GSE8671 and colon cancer cell lines GSE18560 are independent. Thus, except for Ref BN and Setup IV all other setups do not give a consistent indication of significance between the GSE8671 and colon cancer cell lines GSE18560.

The results show that McNemar test is one of the way to test the significance of datasets when the design setups are same as well as different. But hypothesis testing for statistical significance of design setups of the network (with same structure) built on a single dataset might not give clear insight and one needs to be careful regarding this matter. The author currently lacks the awareness of resolving the statistical issue of testing significance of design setups of networks with same structure built on a single data. Finally, the hypothesis testing works well when biological knowledge has been integrated properly into the models as has been shown in Sinha^4^. This work has tried to answer the two questions (1) Are predictions from BNs accurate under different setups? and (2) What is the veracity of obtained results in terms of reproducibility? The first question was answered via generation of accurate results using different training dataset on same test sets. The solution to the second question was found by conducting the statistical significance of the results among the various setups.

## 5 Conclusion

Learned BN on real data is able to accurately predict Wnt activity status as seen in above plots (BN = accurate). Learned BN on simulated data closely matches the BNs trained on real data based on significance evaluation on prediction labels (BN = reproducible). Also, if the probeset measurements are noisy with no discriminative power of whether the gene is active or not, then hiding the observations of genes in some setups, lead to noisy estimation of parameters for genes. This does not happen, if the probeset measurements are not noisy. Also, from a statistical point of view, it was found that, hypothetical testing may not be suitable for studying the behaviour of a network under various criteria, given that the training data for the network remains the same. This gets validated in our experiments that when the various conditions captured via missing observations for different nodes, do not show statistical significance in prediction levels when training dataset is same, but do point to major statistical significance in prediction levels when training on different training dataset. Theoretical analysis of finding a solution for such a scenario would be beneficial in establishing statistical significane of various conditions in the absence of two independent training datasets.

## Conflict of Interest

None.

## Acknowledgement

Thanks to - (1) All anonymous reviewers who have helped in refining this manuscript. (2) Netherlands Bioinformatics Centre (NBIC) for funding the project. (3) Dr. ir. R. H. J. Fastenau (Dean of Faculty of Electrical Engineering Mathematics and Computer Science), Dr. Prof. Ir. K. Ch. A. M. Luyben (Rector Magnificus) and Drs. D. J. van den Berg (President) at Delft University of Technology, for providing support for conducting this work and giving permission to submit the manuscript. (4) Dr. Wim Verhaegh (faculty at Netherlands Bioinformatics Centre and a principal scientist at Molecular Diagnostic Lab in Philips Research) for providing technical details of Naive Bayes model for replicating experiments in Verhaegh *et al*. ^6^. (5) Dr. Marcel J. T. Reinders (scientific director of Bioinformatics Research at Netherlands Bioinformatics Centre and a professor at Delft University of Technology) for reviewing the manuscript. 6) Lastly, the author is indebted to Mr. Prabhat Sinha and Mrs. Rita Sinha for financially supporting this project while the author was on educational and work leave.

